# Cross-species predictive modeling reveals conserved drought responses between maize and sorghum

**DOI:** 10.1101/2022.09.26.509573

**Authors:** Jeremy Pardo, Ching Man Wai, Max Harman, Annie Nguyen, Karl A. Kremling, Cinta Romay, Nicholas Lepak, Taryn L. Bauerle, Edward S. Buckler, Addie M. Thompson, Robert VanBuren

## Abstract

Drought tolerance is a highly complex trait controlled by numerous interconnected pathways with substantial variation within and across plant species. This complexity makes it difficult to distill individual genetic loci underlying tolerance, and to identify core or conserved drought responsive pathways. Here, we collected drought physiology and gene expression datasets across diverse genotypes of the C4 cereals sorghum and maize and searched for signatures defining water deficit responses. Differential gene expression identified few overlapping drought associated genes across sorghum genotypes, but using a predictive modeling approach, we found a shared core drought response across development, genotype, and stress severity. Our model had similar robustness when applied to datasets in maize, reflecting a conserved drought response between sorghum and maize. The top predictors are enriched in functions associated with various abiotic stress responsive pathways as well as core cellular functions. These conserved drought response genes were less likely to contain deleterious mutations than other gene sets, suggesting that core drought responsive genes are under evolutionary and functional constraints. Our findings support a broad evolutionary conservation of drought responses in C4 grasses regardless of innate stress tolerance, which could have important implications for developing climate resilient cereals.

**Significance Statement:** Drought is a complex and variable stress that is difficult to quantify and link to underlying mechanisms both within and across species. Here, we developed a predictive model to classify drought stress responses in sorghum and identify important features that are responsive to water deficit. Our model has high predictive accuracy across development, genotype, and stress severity, and the top features are enriched in genes related to classical stress responses and have functional and evolutionary conservation. We applied this sorghum trained model to maize, and observed similar predictive accuracy of drought responses, supporting transfer learning across plant species. Our findings suggest there are deeply conserved drought responses across C4 grasses that are unrelated to tolerance.

## Introduction

Drought is responsible for billions of US dollars in losses each year, and the impacts of drought are most severe in developing regions of the world where food security is already low (1). Water deficit elicits hundreds to thousands of interconnected molecular pathways in plants, and drought tolerance represents a complex, emergent phenotype that is challenging to breed for or separate into major genetic loci (2, 3). Drought is also a difficult stress to apply and quantify, and plant responses to physiologically relevant drought events in the field are often different from those detected under controlled experiments in growth chamber or greenhouse settings (4, 5). These compounding issues represent major challenges for studying drought stress, but they also present an opportunity to leverage systems level and predictive modeling based approaches to understand complex traits in plants.

C4 grasses dominate natural and agricultural settings, and they have evolved a unique set of adaptations that enable an emergent resilience to drought and other abiotic stresses (6). *Sorghum bicolor* (sorghum) is one of the most stress tolerant and highly productive C4 cereals, and it is an important agricultural commodity grown globally for grain, sugar, and biomass. Sorghum was domesticated in the semi-arid Sudaneese savannah of northeast Africa around 4000 B.C.E (7), and subsequently spread westward across the African steppe and throughout the Indian subcontinent and China. The broad geographic and climatic regions where sorghum was historically cultivated has led to significant diversity and local adaptation among cultivars. While sorghum is generally regarded as a drought tolerant crop, there remains considerable variation for abiotic stress tolerance among different sorghum accessions.

Drought tolerance is a highly complex trait in sorghum and numerous developmental and morpho-physiological traits have been correlated with tolerance. Sorghum cultivars are often classified as either pre-flowering or post-flowering drought tolerant with tolerance at both developmental stages a relative rarity (8). Post-flowering drought tolerance is related to stay-green traits that prevent premature senescence (9, 10). Pre-flowering drought is characterized by more varied responses, and reactive oxygen species scavenging, cuticular wax production, and flowering time regulation are important components of pre-flowering drought tolerance in sorghum (5, 10, 11). Numerous previous studies have examined the transcriptomic response of different sorghum genotypes to both pre-flowering and post-flowering drought tolerance in sorghum (11–13). However, these studies focused on the response of two or a few genotypes, limiting their ability to identify conserved and divergent patterns of expression across the diversity of cultivated sorghum.

The broad genetic diversity of sorghum is captured by the sorghum association panel (SAP), which is composed of 400 temperate breeding lines as well as converted tropical lines that collectively represent the bulk of sorghum diversity (14). Association studies have identified genomic regions linked with drought response in sorghum (15) but unlike expression studies, it is challenging to link specific genes to underlying phenotypes. Here, we compared gene expression across the SAP during a natural drought event and leveraged these data to identify conserved and variable drought responses across sorghum lines.

We hypothesize that a core and deeply conserved drought response operates both within sorghum germplasm and across related species, reflecting ancestral adaptations of C4 grasses. Prior studies have observed commonalities in differentially expressed genes under drought stress across diverse angiosperms, but these studies were limited in sampling, species, or tissue breadth (16, 17). The progenitors of sorghum and *Zea mays* (maize) diverged 11.9 mya, and maize and sorghum still share many similar morphological, biochemical, and genetic traits (18). However, sorghum is more drought tolerant than maize (19), creating an ideal comparative system. The drought responses of sorghum and maize have been compared using only one or a few genotypes (19, 20), but these studies are limited because they fail to account for the broad intraspecific variation present in both species.

Here, we compared interspecific variation and conservation of drought response between maize and sorghum as well as intraspecific variation in both species individually. We generated drought and well-watered expression data across 25 diverse sorghum genotypes and 27 diverse maize genotypes. We also leveraged additional new and public sorghum drought datasets (10–12, 21, 22) to develop a predictive model capable of classifying samples as drought responsive based on gene expression. We dissected the model to identify genes involved in drought response and applied our model to maize to elucidate evolutionary conserved patterns across both species.

## Results

### Climate relevant drought responses across sorghum accessions

Physiologically relevant drought stresses are difficult to simulate in controlled settings, and we sought to capture sorghum responses to a natural drought event in an agricultural setting. East Lansing, Michigan experienced a period of below average precipitation corresponding to a mild drought event between June and early July 2020 (Figure 1a). We collected physiological and RNA samples from 25 diverse sorghum genotypes during this natural stress event and four days later after a heavy rainfall event where plants recovered. We found that the sorghum plants had significantly lower relative water content during the dry period compared to recovery, suggesting the plants were experiencing mild water-deficit stress (Figure 1b). We also found that the sorghum plants had higher instantaneous non-photochemical quenching, as measured by the photosynthesis parameter NPQ_t_, during the dry period (Figure 1c) (23). Non-photochemical quenching increases under drought stress as a mechanism to dissipate excess light energy when photosynthesis is carbon limited as a result of stomatal closure (24). However, despite the drop in NPQ_t_, we did not detect any differences in photosynthetic efficiency (ΦII) or linear electron flow, suggesting that the light reactions of photosynthesis were still proceeding at a high pace (Figure 1d). Together, this suggests the sorghum plants were experiencing a very mild, and fully recoverable drought event.

**Figure 1.**
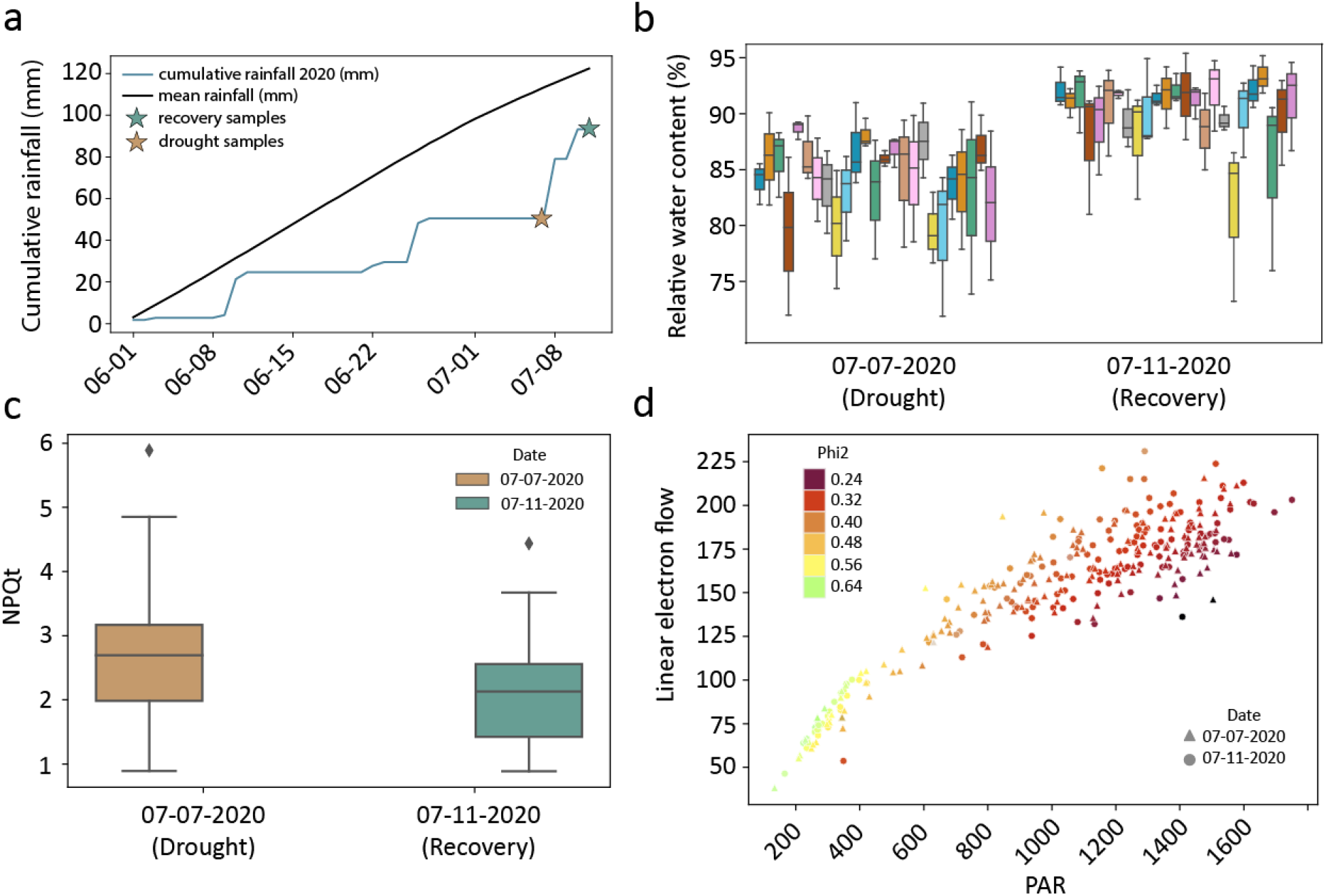
Physiological response of diverse field grown sorghum genotypes across a natural drought stress event. (a). Cumulative growing season precipitation before and during the sampling period compared to the 30 year mean. The two sampling dates are labeled with stars. (b). Box plots of relative water content for each of the 25 genotypes on the two sampling dates. (c) Boxplot of NPQt on each sampling date. (d) Scatterplot showing linear electron flow (LEF) as a function of photosynthetically active radiation (PAR). Points are colored by photosystem II efficiency (Phi2) with circles representing the recovery (7/11/20) sampling date and triangles representing the drought (7/7/20) sampling date.

We collected three replicates of RNAseq data for each genotype at the drought and recovery timepoints to search for expression patterns corresponding to water deficit responses in sorghum. Dimensionality reduction analysis clearly separates the RNAseq samples into two distinct groups of well-watered and drought along principal component 1 (Figure 2a). Across all genotypes, we identified 1,761 genes upregulated under water deficit, and 2,317 genes downregulated (Supplemental table 1). Among upregulated genes under drought, we found enrichment of gene ontology terms related to stress responses, including response to heat as well as terms related to protein folding and chaperone activity (Supplemental Table 2). Genes downregulated under stress were enriched in gene ontology terms related to photosynthesis and central metabolism, as expected for mild stress responses (Supplemental Table 3).

**Figure 2.**
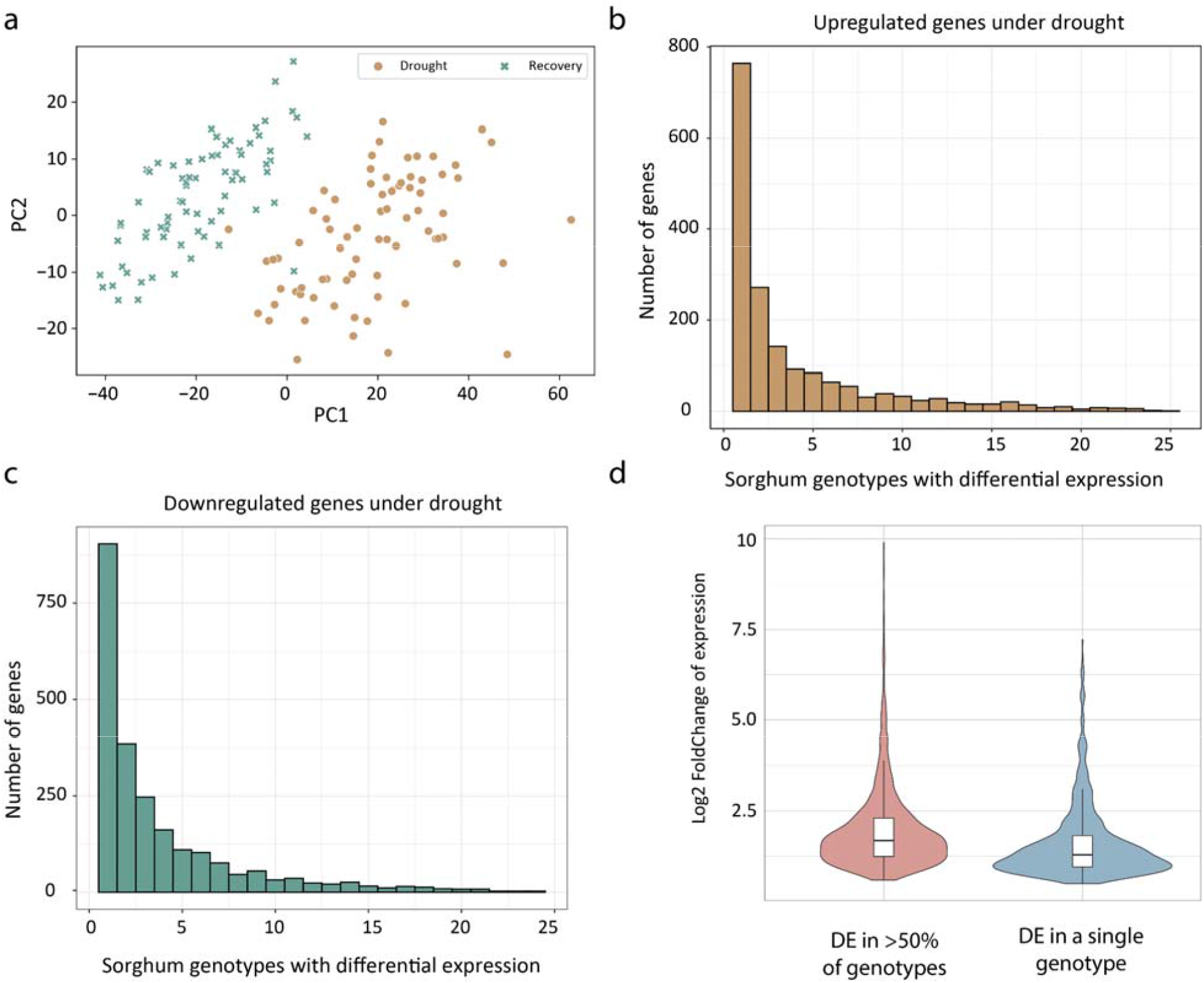
Unique expression signatures of drought stress across sorghum genotypes. (a) Principal component analysis of log2 transformed RNAseq data for the sorghum field drought experiment. Individual samples are plotted and colored by day. (b) Histogram showing the number of shared upregulated expressed genes across the 25 sorghum accessions. (c) Histogram showing the number of shared downregulated expressed genes across the 25 sorghum accessions. (d) Violin plots of log2 fold change of expression in the shared differentially expressed genes compared to the genes uniquely differentially expressed in a single genotype.

Despite the large overall changes in gene expression and typical stress-related gene ontology profile, we found surprisingly little intraspecific overlap of gene expression under water-deficit stress in sorghum (Figure 2b, c). Only a single sorghum gene was upregulated and no genes downregulated in all 25 genotypes on the drought sampling date compared with recovery. We defined a set of 269 “shared” differentially expressed genes based on common differential expression between the drought and recovery timepoints in at least half of the sorghum genotypes. Only 133 genes, representing 8% of all upregulated genes, showed shared upregulation. Similarly, only 136 genes or 6% of downregulated genes were shared in half or more genotypes. On a per genotype basis, a greater percentage of the differential expressed genes were shared, with between 18% and 51% of genes differentially expressed in a given genotype being shared. The mean percentage of upregulated genes in each genotype that were shared across at least half the genotypes (36%) was significantly higher (t-test p = 0.01) than the percentage of shared downregulated genes (28%). To further explore the difference between shared and variably expressed genes, we defined a set of 1,583 “unique genes” which were differentially expressed in only a single genotype. While the absolute number of unique genes is higher than the shared genes overall, in any given genotype they represent a lower percentage of the up and downregulated genes. We found that the log2 fold-change for shared upregulated genes was significantly higher as compared with unique upregulated genes (Figure 2d; t-test p=1.48e-15). We compared gene ontology enrichment between the set of shared upregulated and unique upregulated genes to ascertain possible differences in function between the two sets. Gene ontology enrichment of the shared upregulated genes mirrored that of all upregulated genes, with terms related to response to heat, protein folding, and reactive oxygen species scavenging enriched. Conversely, gene ontology terms enriched among unique upregulated genes were less obviously related to stress response with terms such as translation, peptide metabolic process, and cellular amide metabolic process enriched.

### Predictive modeling of drought responsive genes in sorghum

Our differential expression analysis was limited by either the relatively mild nature of the natural drought event and/or the low number of replicates per genotype (three), potentially causing us to miss shared drought responsive genes due to insufficient statistical power. We used a predictive modeling approach to more robustly identify any shared water-deficit stress responses across sorghum genotypes in our experiment. Our approach involved training a random forest model to classify samples as “drought” or “control” based on normalized gene expression values alone. We hypothesized that the features with the most predictive power in the model would represent genes with central and conserved roles in drought responses. We first applied this approach to the sorghum experiment described above. Using a training set of expression from 75% of sorghum genotypes, our model had near perfect prediction accuracy on the remaining test set across all folds in a five-fold cross validation scheme. However, the model relied on a small number of features to make those predictions. The average depth of the individual trees within the random forest was only 1.9, indicating that each decision tree used on average less than two genes out of 34,117 possible genes to make a prediction. To improve the utility of our model, we used k-means clustering to reduce the total number of features. We created seven clusters based on the scaled expression data and used the first principal component of gene expression in each cluster as our input feature. Our model was able to classify samples from genotypes withheld from the training data correctly 95% of the time (Supplemental Figure 1). The individual trees within the cluster-based model used on average 4.8 out of seven possible k-means based features. To identify clusters with the most predictive power, we calculated feature importance using the mean decrease in impurity (GINI score) metric implemented in the scikit-learn package. We found that clusters with the most importance in the classification model also had the highest percentage of upregulated genes (Supplemental Figure 2).

Our model was developed using data from a relatively mild drought event, and we expanded the model using publicly available sorghum drought datasets with varying designs, genotypes, and drought severity (Supplemental Table 4). The public datasets included RNAseq of vegetative tissues from both field and chamber grown sorghum across multiple developmental stages. We also generated an additional dataset using 54 chamber grown Btx623 sorghum plants with drought applied at three different developmental stages. In total, we analyzed seven additional datasets, collectively representing 35 genotypes, with 206 drought stressed samples and 254 well-watered or recovery samples. We reprocessed all of the expression data using a common analytical framework, and compared these experiments using dimensionality reduction approaches. The expression samples cluster separately by experiment rather than stress vs control along the first two principal components, suggesting significant heterogeneity and sampling artifacts (Figure 3a). To remove batch effects, we applied the combat algorithm to adjust the input data (Supplemental Figure 3). We then split the data into training and testing sets using a “leave one experiment out” approach where one dataset was withheld for testing the model and the rest were used for training. The accuracy of our model across all test datasets was 86% (Figure 3b). The precision and recall of the model were both 0.84, where 1 is a perfect classifier and 0.5 is a random classifier (Figure 3c, d). The model performed well across all datasets individually, with prediction accuracy ranging from 64% to 100%, however excluding the best performing (which only had two samples) and worst performing datasets, the range was 82% to 91% (Figure 3b). The ability of our model to classify samples accurately on an unobserved dataset implies the existence of a conserved pattern of gene expression in response to drought across diverse sorghum lines.

**Figure 3.**
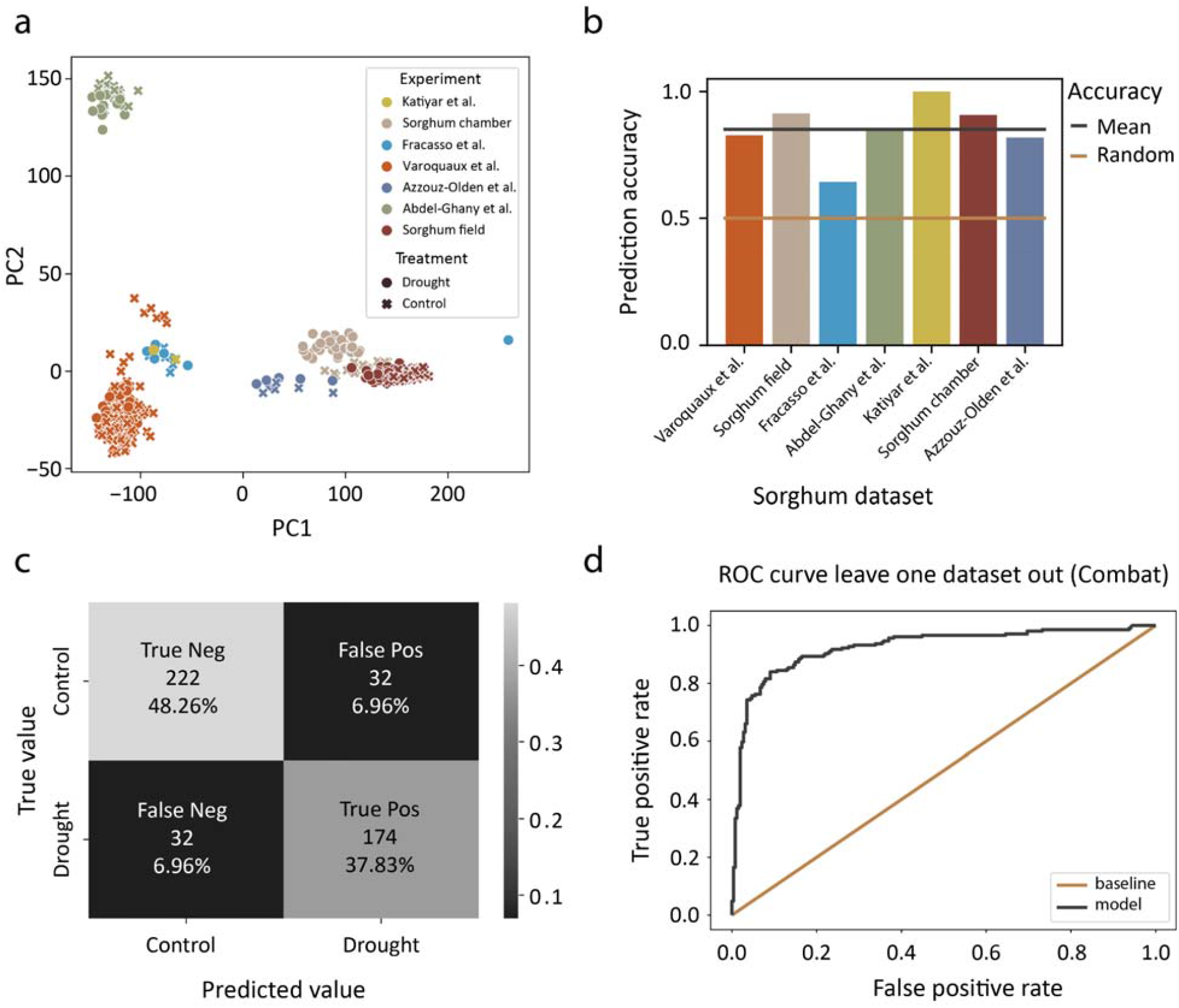
Predictive modeling of drought stress in sorghum using gene expression data. (a) Principal component analysis of log2 transformed RNAseq data for the seven sorghum drought expression datasets used for predictive modeling. A PCA of the ComBat-filtered expression data is available in Supplemental Figure 3. (b) Predictive accuracy of the random forest model for classifying drought stressed sorghum samples across each individual experiment. The mean predictive accuracy is shown by a black line compared to a random background (in orange). (c) Confusion matrix of the drought predictive model. (d) Receiver operating characteristic curve showing the performance of the drought classification model across all classification thresholds.

Developmental stage influences the relative drought tolerance of sorghum lines. To assess whether our model could accurately classify drought samples regardless of developmental stage we trained a version of our model using 203 sorghum samples from the vegetative stage and used the remaining 34 samples from flowering or post-flowering stages to test the model. Our model predicted with 97% accuracy and an auc score of 0.99 (Supplementary Figure 4), suggesting that a conserved drought response is present across developmental stages, despite distinct physiological signatures and molecular mechanisms underlying pre and post flowering drought responses in sorghum.

**Figure 4.**
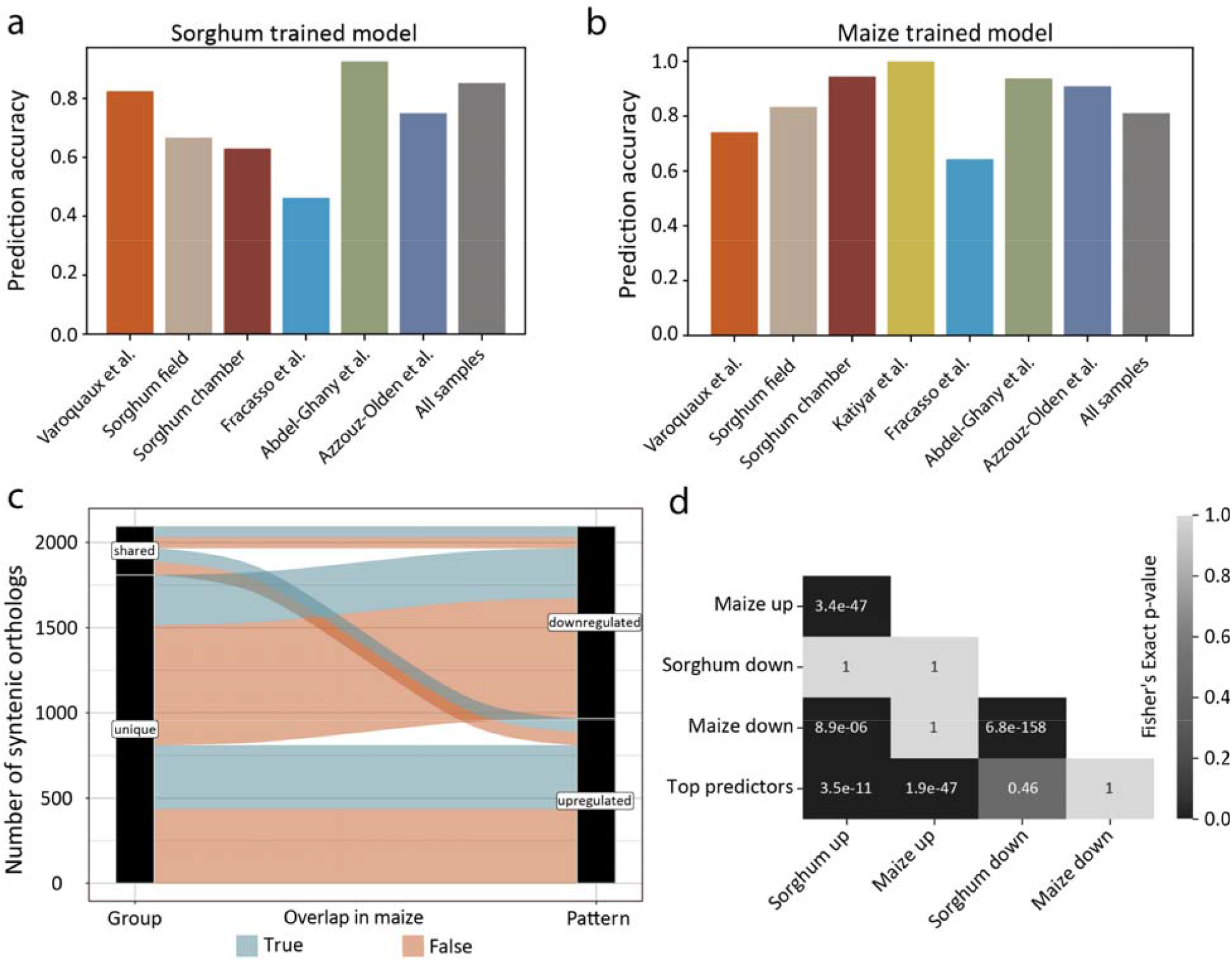
Cross species predictive modeling of drought stress. (a) Predictive accuracy for classifying drought stress in maize using all of the sorghum samples for training (in gray) and each experiment individually. (b) Predictive accuracy of the maize trained model for classifying drought stress across the sorghum experiments. (c) Alluvial plot showing orthologs between maize and sorghum that are conserved top predictors (blue). (d) p-value from Fisher’s exact test comparing overlap between syntenic orthologs in each differentially expressed gene set as well as the top predictors from the sorghum trained model.

### Cross-species predictive modeling identifies conserved core stress response

Our analyses to this point identified a shared drought response across diverse sorghum lines. To probe the evolutionary conservation of this response, we compared our findings within sorghum to similar datasets in maize. We collected a water-deficit stress dataset across a set of 27 diverse maize genotypes in a greenhouse environment. Briefly, we withheld water from potted maize plants at the ∼V5 leaf stage for one to three days. On each day, we sampled a stressed group and a corresponding control group, which received water daily. We found that stomatal conductance was significantly lower in the stressed groups as compared with the controls (p=1.22e-105) as well as across the different experimental timepoints (p=1.04e-31) (Supplemental Figure 5). As expected, the difference between days was dependent on treatment with a significant interaction between treatment and day (p=1.43e-38), demonstrating that the stressed group had a significant drop in stomatal conductance compared with the controls. We also found significant differences in stomatal conductance between genotypes (p=0.0017) suggesting differing physiological responses across maize genotypes.

**Figure 5.**
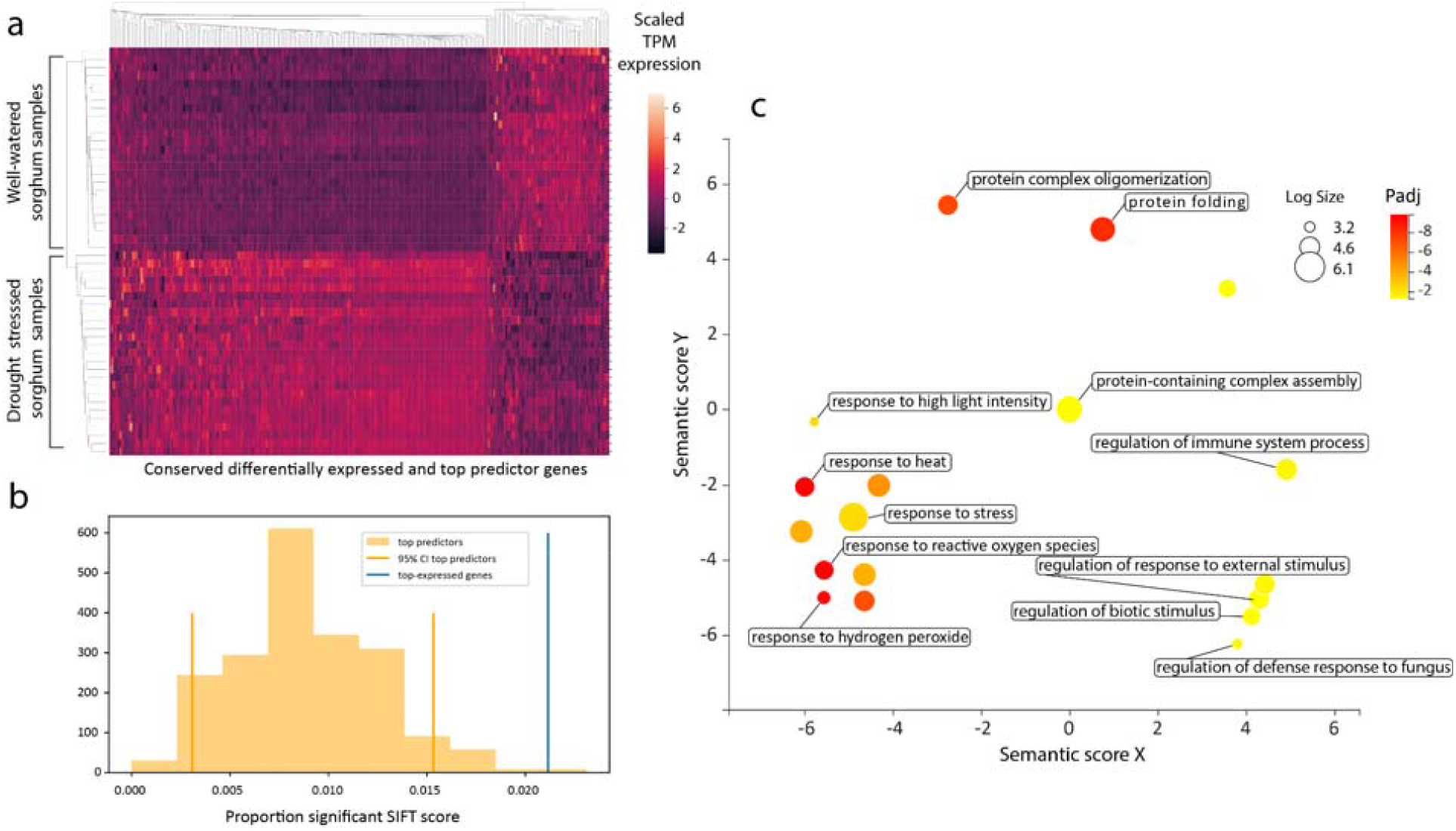
Evolutionary conservation and functional enrichment of top predictors involved in drought responses. (a) Heatmap showing scaled expression values in the sorghum field experiment, for the 284 syntenic orthologs that are differentially expressed in both the maize and sorghum experiments as well as among the top-predictors in the sorghum and maize trained models. (b) Bootstrapped confidence interval for the proportion of genes with a significant average SIFT score among the sorghum trained model top predictors compared with the proportion of top expressed genes (>74 percentile of expression) with significant average SIFT scores. (c) Multi-dimensional scaling plot showing clusters of enriched gene ontology terms in the set of genes described above. The size of each circle is proportional to the number of genes annotated with each term and the circles are colored by the log10 of the adjusted p-value.

We observed significant changes in gene expression between the drought and control timepoints in maize, with the number of differentially expressed genes increasing in the more severe timepoint. The maize dataset has only one sample of each genotype at each timepoint, thus instead of comparing differential expression for each genotype, we used log2 fold-change to assess the variation of expression response under water-deficit across genotypes. Similar to the sorghum dataset, we saw limited shared response between the genotypes. Only three genes had a log2 fold-change greater than 1.5 between the first, milder, stress timepoint and the corresponding control in all 27 genotypes. While 272 genes, representing 7% of all upregulated genes showed a greater than 1.5 fold-change in at least half the genotypes. During the severe stress timepoint, we saw more overlap between genotypes with 125 genes showing a greater than 1.5 fold-change across all genotypes and 1,845 upregulated in at least half of the genotypes.

We used our modeling approach to test whether sorghum and maize have shared, core drought response pathways. We converted maize genes to their corresponding sorghum orthologs using a synteny based approach to enable comparisons across species. Although both sorghum and maize share the same chromosome number, maize underwent a more recent whole genome duplication and many maize genes display a 2:1 syntenic pattern with sorghum (Swigoňová et al., 2004). For maize genes with this 2:1 synteny pattern, we averaged syntelog expression and created a converted matrix of maize expression with sorghum gene identifiers. We then retrained our sorghum model using all 7 sorghum datasets with only sorghum genes having a syntenic counterpart in maize (Figure 4a). We applied the model to the maize data and found that it predicted with 85% accuracy and an auc score of 0.98. We also created a new model trained with the maize data, and tested it on the sorghum dataset. This maize model predicted sorghum samples with 81% accuracy and an auc score of 0.94 across all samples with performances of 64%-100% across the individual experiments (Figure 4b).

To further test the hypothesis that maize and sorghum share a core stress response, we compared the overlap between differentially expressed genes in the maize dataset, our sorghum field experiment, and the top-predictors from our sorghum trained model. We found significant overlap between genes upregulated in maize and sorghum (fisher’s exact test p = 3.4e-47), as well as genes downregulated in the two species (fisher’s exact p = 6.8e-158; Figures 4c, d). We also found significant overlap between the top predictors from the sorghum trained model and genes upregulated in maize (fisher’s exact p = 1.9e-47) as well as sorghum (fisher’s exact p= 3.5e-111). Interestingly, we did not identify significant overlap between downregulated genes and the top predictors in our model (Figure 4d).

We hypothesized that expression of the top predictors from our sorghum trained model would be associated with physiological markers of drought stress. To test this, we calculated the first principal component of log transformed gene expression (PC1) across the top predictors as a summary value of gene expression (Supplemental Figure 6). We then correlated the PC1 values with physiological variables. We found that PC1 was significantly negatively correlated with relative water content (spearman r = −0.53), suggesting that the top predictors are strongly associated with signatures of drought physiology.

We identified a set of 284 genes that were top predictors in the sorghum and maize trained models, and also differentially expressed in both datasets. The majority of these genes showed increased expression during drought in our sorghum dataset (Figure 5a). Using gene ontology enrichment analysis, we found significant enrichment for well-characterized abiotic and biotic stress responsive pathways as well as genes related to protein folding (Figure 5c). We found that these conserved drought responsive genes were also significantly more likely to have shared differential expression (>50% of genotypes) as opposed to differentially expressed in only one sorghum genotype (fisher’s exact p = 7.37e-29). Previous researchers have identified sets of shared differentially expressed genes related to drought responses in other species (17). To test for overlap between our conserved drought genes in maize and sorghum and across broader species, we used conserved orthogroups to link gene identifies between studies. We identified orthologs for 282 of the 284 conserved drought responsive genes reported in Shaar-Moshe et al. and found 39 had shared drought responsiveness in maize and sorghum. This represents significant enrichment (fisher’s exact test p = 2.49 e-20), however unsurprisingly a substantial portion of the shared response genes between maize and sorghum are not shared with more distantly related species.

### Evolutionary constraint of conserved drought responsive genes

Through our predictive modeling approach, we have identified a core set of genes that show a conserved pattern of gene expression during water deficit in maize and sorghum. We expect that the shared expression signatures are an indication of evolutionary conservation. To test this, we compared deleterious load between top predictor genes and a background set of genes. To estimate deleterious load we used average SIFT scores as calculated in Lozano et. al. 2021 (25). SIFT scores are computational predictions of the effect of individual mutations. A SIFT score below 0.05 represents a mutation that is predicted to be deleterious, and when averaged across all mutations in a gene the score represents a deleterious index. We compared the proportion of genes with average SIFT scores below 0.05 between 2000 bootstrapped samples of the top predictor genes with a background set. Since genes with high expression are more likely to be evolutionary constrained, we chose the set of all genes with average expression values across all our sorghum field samples greater than the 73rd percentile, which represents the mean percentile rank of the top predictor genes. The top predictor genes had a significantly lower proportion of genes with average SIFT scores < 0.05 than both the highly expressed background set and all genes (Figure 5b). This suggests that the core set of drought related genes are more evolutionarily constrained than other highly expressed genes.

## Discussion

Drought tolerance is variable across diverse sorghum lines, yet some elements of drought response are conserved even across species. Previous work has mostly focused on either differences between individual sorghum genotypes or comparisons of sorghum with other species such as maize. Integrating our understanding of intraspecific and interspecific variation in drought response is an important step in unraveling the evolutionary history of drought tolerance in plants. In this study, we used a predictive modeling approach combined with differential expression analysis across diverse sorghum genotypes to identify shared and unique drought responses.

We identified a core set of genes with a conserved expression pattern across the majority of sorghum genotypes. We then applied our model to a parallel maize dataset and found that the conserved response was largely shared with maize. In evolutionary terms, the ancestors of maize and sorghum diverged relatively recently (18). The two species show conserved response to some stresses, and previous studies have shown conserved resistance mechanisms to particular pathogens between maize and sorghum (26). However, sorghum and maize differ markedly in their resilience to abiotic stresses, particularly drought and heat (19, 27, 28). Interestingly, even for cold stress, an abiotic stress where both species are susceptible, maize and sorghum have surprisingly different gene regulatory responses (29). Therefore, our finding that a core response to drought is conserved between maize and sorghum is initially surprising. A meta-analysis of microarray data identified shared differentially expressed genes across multiple angiosperm species during progressive drought stress, although this work did not include sorghum or maize (17). Our findings expand on this result, showing a similar pattern of conservation in sorghum and maize during drought. Core aspects of angiosperm drought response evolved during the adaptation of early plants to a terrestrial environment (30). Conversely, cold tolerance likely evolved repeatedly across angiosperm lineages and relatively recently in grasses (31). The apparent divergent responses to cold in sorghum and maize and seemingly more shared drought response are perhaps an artifact of the evolution of these two traits across different timescales.

While prior work used differential gene expression to identify shared patterns across species, we used a combination of differential gene expression and a predictive modeling approach. Small sample sizes can limit the effectiveness of differential gene expression analysis (32). Combining samples from disparate datasets can increase the sample size, however, this is impractical due to differences in methods between experiments. In particular, drought experiments often represent a broad range of soil water contents, developmental stages and genotypes.

Supervised classification models offer an alternative approach to traditional differential gene expression analysis. We used a random forest classifier to label samples as “drought” or “control” based on gene expression values. The random forest model was able to accept training data from seven diverse datasets which varied in sample size, growth environment, developmental stage, and method and level of water-stress imposed. Our model performed well across the majority of these datasets indicating a broadly shared core drought response across disparate sorghum drought datasets. Developmental stage has a major impact on drought tolerance in sorghum, and separate pre or post flowering drought tolerant accessions have been identified, with little overlap between groups (5, 9). Pre and post flowering drought tolerance strategies are characterized by distinct physiological and molecular mechanisms. Post flowering tolerance is associated with the stay green phenotype where tolerant lines retain green leaf area from anthesis through grain filling. The physiological basis of pre flowering drought tolerance is more complex, and likely relates to water use efficiency, osmotic adjustment, and plant architecture traits that ultimately give rise to higher yield (33). Despite these differences, when trained on only the vegetative stage samples, our model still classified flowering and post-flowering drought samples accurately. This implies a shared core stress response across developmental stages.

Despite broad conservation of a core set of drought responsive genes across sorghum datasets and developmental stages, the individual expression response to drought was variable across sorghum genotypes. The majority of differentially expressed genes identified in our sorghum field experiment were private to one genotype. However, within a single genotype on average a higher percentage of upregulated genes are shared (36%) than genes which are unique to that genotype (8%). The private genes are potentially responsible for between genotype differences in drought response. Alternatively, they may represent noise or gene expression changes unrelated to drought. Overall, the log2 fold-change of shared genes was significantly higher than unique genes, suggesting that unique genes are more likely to represent noise rather than true differentially expression. Furthermore, gene ontology terms enriched among unique genes were not clearly stress related while the shared genes were enriched in terms related to known stress response pathways. While some unique differentially expressed genes are undoubtedly important in drought response, we hypothesize that the core drought response is conserved across genotypes.

Our finding of a conserved drought response across diverse sorghum genotypes and developmental stages coupled with the cross-species predictive accuracy of the sorghum and maize trained models suggests an evolutionarily conserved response. A prior meta-analysis found that differentially expressed orthologs which were shared between wheat and rice or barley and rice had higher sequence similarity than orthologs which were differentially expressed in only one species (17). Not all sequence changes are functionally meaningful. Top predictors in our sorghum-trained model had a significantly lower proportion of genes with average SIFT scores below 0.05 (i.e., predictive of deleterious mutations (34)) than either a random set of background genes or other highly expressed genes. This suggests that conserved drought responsive genes across sorghum and maize are less likely to contain deleterious mutations.

Several metabolic processes have repeatedly been shown to be involved in drought response across divergent plant species. Gene ontology terms related to response to abiotic stimulus and carbohydrate metabolism were identified as enriched among conserved differentially expressed genes in Shaar-Moshe et. al. Other studies proposed that pathways involved in accumulation of osmoprotectants, reactive oxygen species scavenging, regulation of nitrogen metabolism, ammonia detoxification, and activation of the GABA shunt in the TCA cycle were conserved across multiple species in response to drought (16). The core sorghum and maize responsive genes we identified have overlap between orthogroups identified in Sharr-Moshe et. al. and the general gene ontology term “response to abiotic stimulus”. We also found evidence of reactive oxygen species scavenging enzymes as well as folding and refolding of proteins based on the GO term enrichment. Cellular response to endoplasmic reticulum stress caused by accumulation of unfolded and misfolded proteins, known as the unfolded protein response (UPR) is a well-studied process in response to environmental stress (35). Much of the UPR is conserved across not just plants but all eukaryotes and thus it is unsurprising that we see shared activation under drought stress here (36).

## Conclusion

Prior studies have identified shared differentially expressed genes under drought stress across multiple species. We extend their results showing a similar shared core response across maize and sorghum using a novel predictive modeling approach. Our approach has the advantage of enabling integration of multiple diverse datasets despite differences in sample size and approach between experiments. We show that the core response is largely shared among diverse sorghum genotypes and across developmental stages despite overall variable drought response between species. Taken together, our results suggest a deeply conserved core drought response modified by individual variation.

## Methods

### Sorghum experimental design and sampling

We grew *Sorghum bicolor* for this experiment at the Michigan State University Agronomy farm using a randomized complete block design. The soil type was a mix of Conover loam over approximately two thirds of the field area, and the more freely draining Sisson fine sandy loam, in the remaining area (U.S. Department of Agriculture, Natural Resources Conservation Service, 2019). We planted seeds in two row plots and allowed the plants to grow under ambient environmental conditions. East Lansing, Michigan experienced a drier than normal period during the early summer of 2020. The nearby Hancock Turf Research Center weather station recorded only 50.5 millimeters of precipitation between June 1st and our first sampling date of July 7th compared to the 5 year average of 106.68 millimeters at that site. The end of June and beginning of July was particularly dry with no precipitation falling between June 27th and the first sampling date of July 7th. In total, 42.6 mm of rainfall fell before the second sampling timepoint on July 11th.

We sampled each plot at two separate timepoints, the first on July 7th, 2020, was during mild water-deficit stress, and the second on July 11th was after the plants had recovered following precipitation. On both days, we took all samples between 10:00 am and 12:00pm local time and sky conditions were similar. We collected leaf samples for RNA sequencing into liquid nitrogen from the midsection of the second top-most fully expanded leaf from three plants per plot and combined the samples into a single tube. Leaf tissue from the same leaves were collected into airtight tubes and stored in a cooler for relative water content analysis. We also collected photosynthetic efficiency and other leaf physiology data using the MultiSpeQ fluorometer from the top-most fully expanded leaf for two plants per plot.

We measured the fresh weight (FW) of leaf samples using an analytical balance immediately following field sample collection. We processed three leaf samples from each plot together to achieve a single relative water content value per plot. After measuring fresh weight, we floated the leaf samples in Millipore filtered deionized water kept in the dark overnight at 4°C. The following day, we dried the surface of the leaf samples and measured the turgid weight (TW) and placed the samples in paper envelopes to dry at 60°C. After drying overnight, we measured the sample dry weight (DW) and calculated relative water content using the formula:

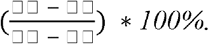

### Maize experimental conditions and sampling

For the maize drought experiment, we grew the 26 founders of the NAM population, as well as the inbred maize line Mo17, in 4” diameter by 4” deep nursery pots for three weeks during the month of June 2016 in the Gutterman greenhouse located in Ithaca, NY (42.4482 N, 76.4612 W) (37). Supplemental lighting in the greenhouse provided a minimum of 300 μmol M^−2^ S^−1^ of PAR. During strong sunlight, PAR typically approached 1000 μmol M^−2^ S^−1^. The temperature in the greenhouse was held at approximately 28°C during the day and 20°C at night. We hand-watered plants twice daily except during drought treatments. We seperated the plants into three blocks in the greenhouse with each block consisting of one complete set of genotypes for the control and drought treatments. After three weeks of growth, at approximately 5th leaf stage, we withheld water from the drought treatment. We measured stomatal conductance on each day of the experiment beginning on day 0 (both the control and drought treatments were well watered) and ending on day 3 (three days after cessation of water for the drought treatment). A Decagon SC-1 porometer, calibrated each day of the experiment, was used to collect all stomatal conductance readings from the uppermost fully expanded leaf. All stomatal conductance measurements were collected between 10:00 AM and 2:00 PM EDT to minimize the impact of daily physiological cycles on the readings. Pots within the drought treatment were weighed each day of the experiment as a proxy measure for soil moisture. On days 1 and 3 of the experiment, we collected leaf tissue for RNA sequencing in liquid nitrogen. We selected the second top most fully expanded leaf or tissue collection to avoid competition with the leaves selected from physiological measurement. We sampled tissue by folding the leaf from the tip to the base and excising an ∼5 cm section spanning the midpoint of the leaf and extending inward to, but not including the midrib. On day 3, samples were collected from the other half of the second topmost fully expanded leaf when possible. However, in cases where the prior sampling had damaged the leaf, the topmost fully expanded leaf was used as a replacement.

### RNAseq Profiling

For both the sorghum and maize experiments, we excised a leaf section from the midpoint of the second top-most fully expanded leaf from three plants per plot (sorghum experiment) or pooled samples from three plants (maize experiment) and froze them in liquid nitrogen. We lysed frozen leaves using a bead tissue homogenizer. We then thawed the ground tissue in trizol reagent and extracted RNA using a Direct-zol 96 kit according to manufacturer’s instructions (Zymo Research, Irvine CA). Lexogen quant-seq libraries for each sample were prepared and sequenced by the Cornell Institute of Biotechnology for the maize and sorghum datasets.

Previously published RNAseq data for drought stress in sorghum was collected from (10–12, 21, 22) and downloaded from the NCBI sequence read archive and processed as described below. Full details of the published RNAseq data can be found in Supplemental Table 4.

### RNA Sequence processing

We trimmed sequence adapters and quality checked the raw FASTQ files using the program fastp (v0.23.2) (38). We then pseudo-aligned our cleaned sequencing reads to the Btx623 sorghum or B73 V5 maize reference genomes using salmon (v1.6.) (39–41). We then converted transcript level counts to gene level using the R package TXimport (v 1.22.0) (42). We used DESeq2 (v1.36.0) to calculate pairwise differential expression between drought and well-watered conditions for each genotype (43).

### RNAseq data normalization and batch effect removal

For our model built across sorghum datasets, we removed batch effects using the combat algorithm implemented in the python package pyComBat (v0.3.2) (44). For all models, we split the data into training and testing sets using approaches outlined in Supplemental Table 5. After splitting the data into training and test sets we scaled the data using the Standard scalar function from the scikit-learn python package (v1.1.0) (45).

### Random Forest Model Construction and feature importance

We constructed random forest models with the RandomForestClassifier function from scikit-learn (v1.1.0) (45). To select hyper-parameters, we used the RandomizedGridSearchCV function with 100 iterations using 3-fold cross-validation to search the parameter space (Supplementary Table 6).

We calculated feature importance using mean decrease in impurity (Gini score) as implemented in the scikit-learn package (v1.1.0). We then ranked all genes by their importance score. To identify a set number of “top predictors” we used a heuristic approach whereby we selected the *n* top features and compared the number that overlapped with differentially expressed genes with the number of overlaps in a random set of *n* genes. For each set of size *n* we calculated a z-score (# of *n* top predictors that are also differentially expressed – mean (# of *n* randomly selected genes that are differentially expressed) / standard deviation of random genes. We then selected 675 top predictor genes, as that maximized the z-score.

## Data availability

RNAseq data generated in this project are available on the NCBI sequence read archive for maize and sorghum.

## Acknowledgements

This work is supported by NSF Grant MCB□1817347 (to R.V.). M.H was a participant in the Plant Genomics Research Experience for Undergraduates Program funded by NSF BIO Division of Biological Infrastructure (NSF-DBI 1358474). J.P. was supported by predoctoral training award T32-GM110523 from the National Institute of General Medical Sciences of the NIH.The authors declare no competing financial interests.

## Notes

### Competing Interest Statement

The authors have declared no competing interest.

