## Supplemental Figures/Tables for "Cross-species predictive modeling reveals conserved drought responses between maize and sorghum"

**Supplemental Figures and Tables**

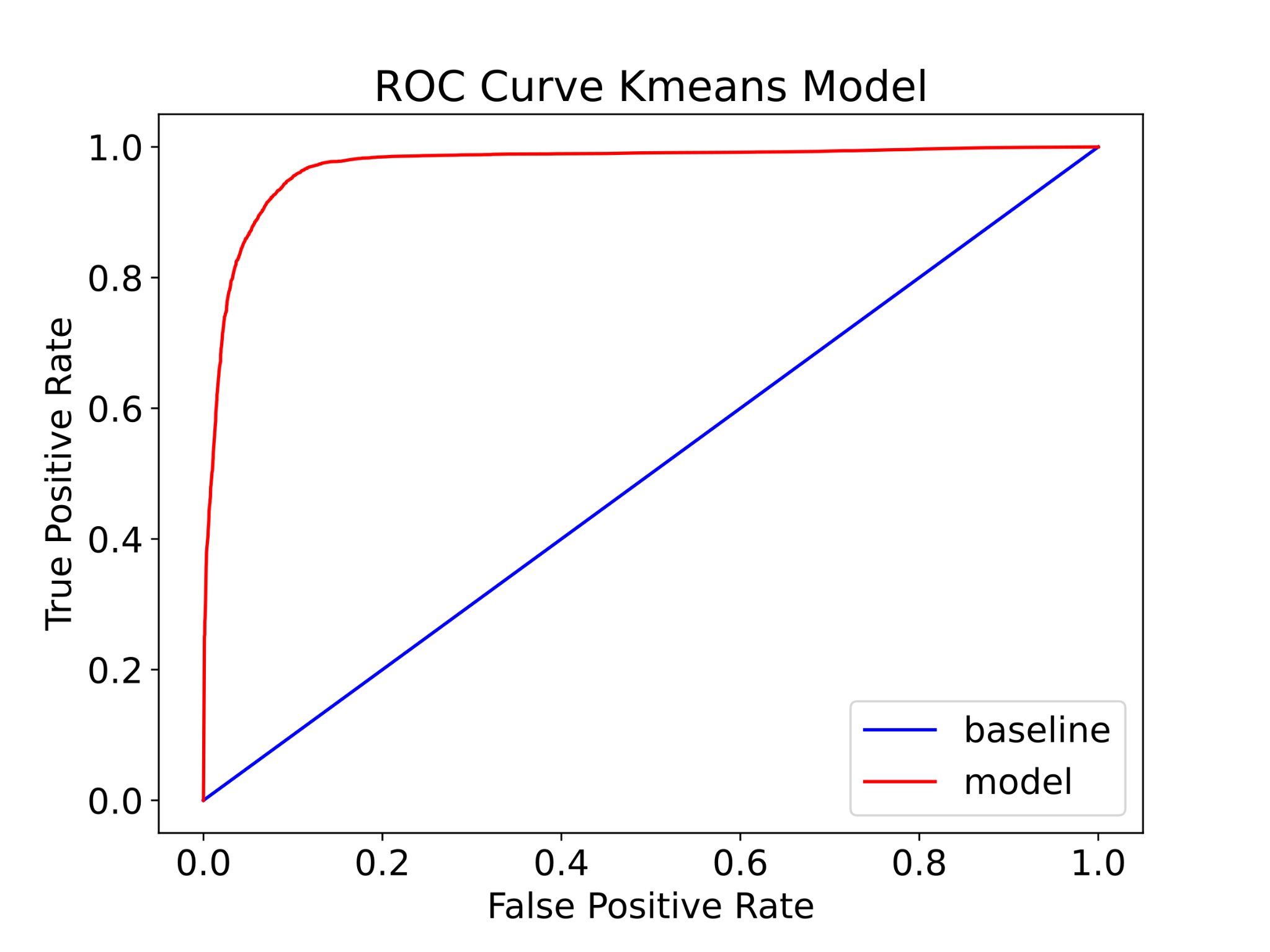

**Supplemental Figure 1. AUC-ROC curve for 500 replications of the random forest model using the kmeans clusters.**

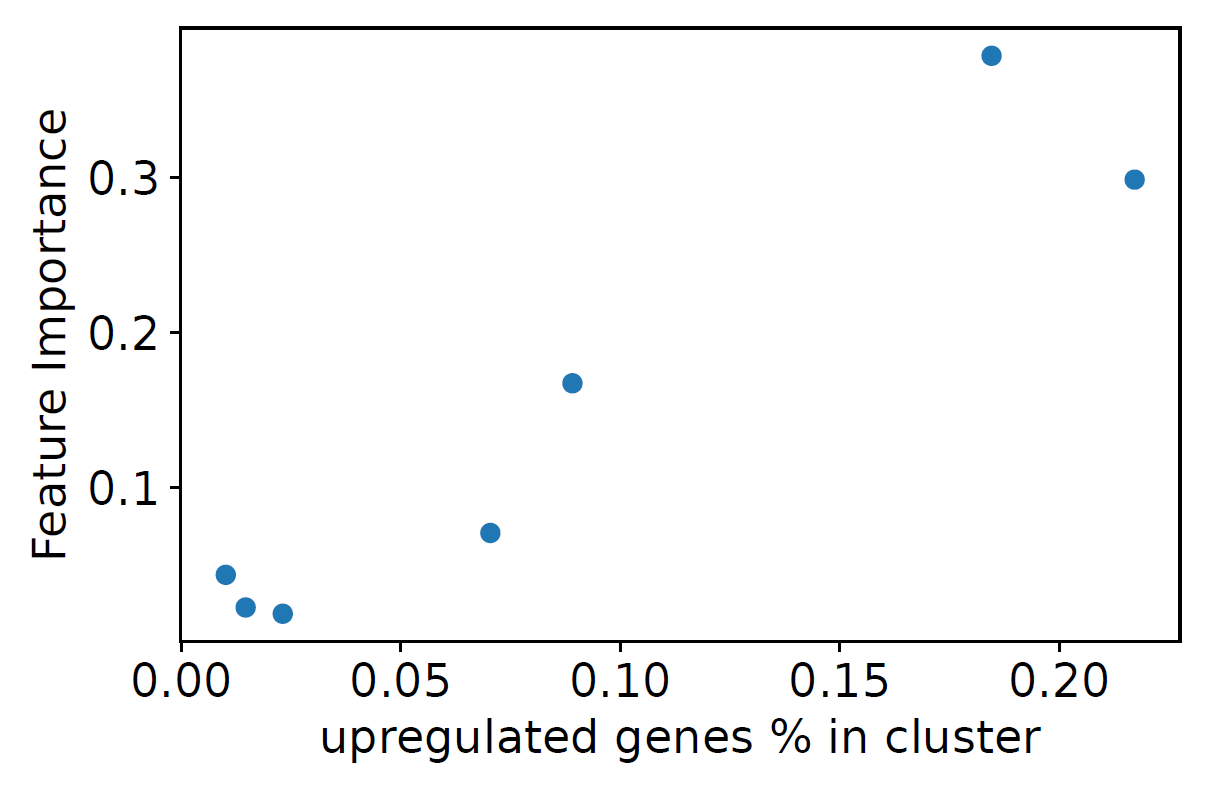

**Supplemental Figure 2. Feature importance for the random forest model using the kmeans clusters.** The proportion of upregulated genes in each kmeans cluster compared to feature importance is plotted.

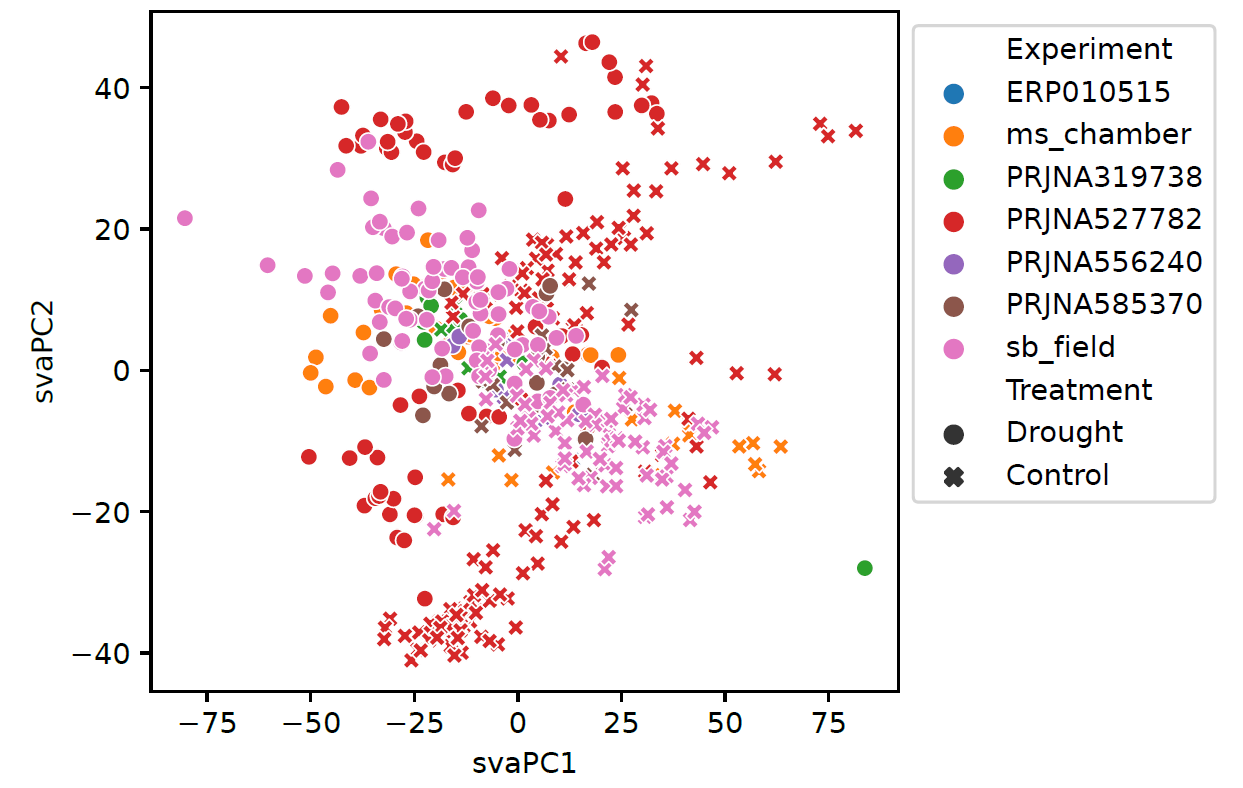

**Supplemental Figure 3. Principal component analysis of ComBat adjusted sorghum RNAseq data.** Each of the seven sorghum experiments is shown by color with drought samples shown as circles and control (or well-watered) as crosses.

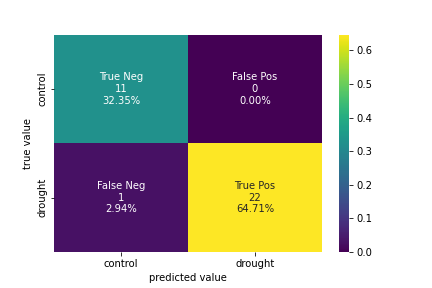

**Supplementary Figure 4. Confusion matrix showing model performance for classifying flowering and post-flowering stage samples using a classifier trained using 203 vegetative stage samples.**

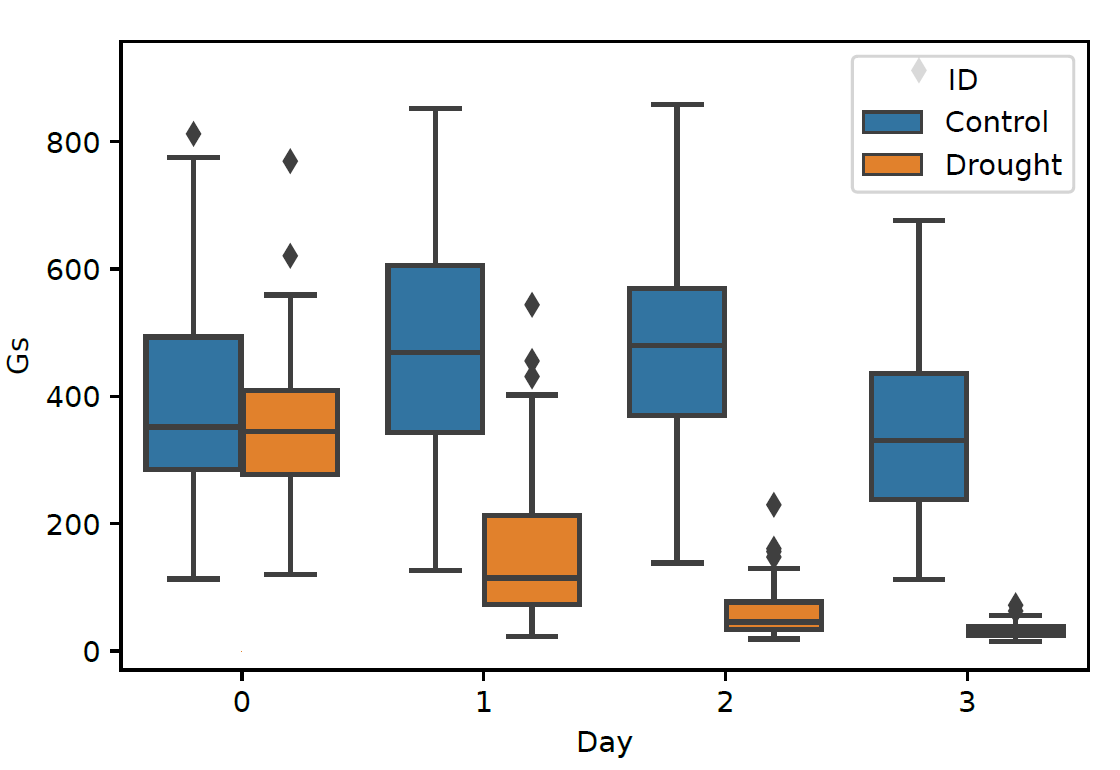

**Supplemental Figure 5. Gas exchange for the 27 NAM founder maize lines for the control (blue) and drought (orange) samples throughout the drought timecourse.**

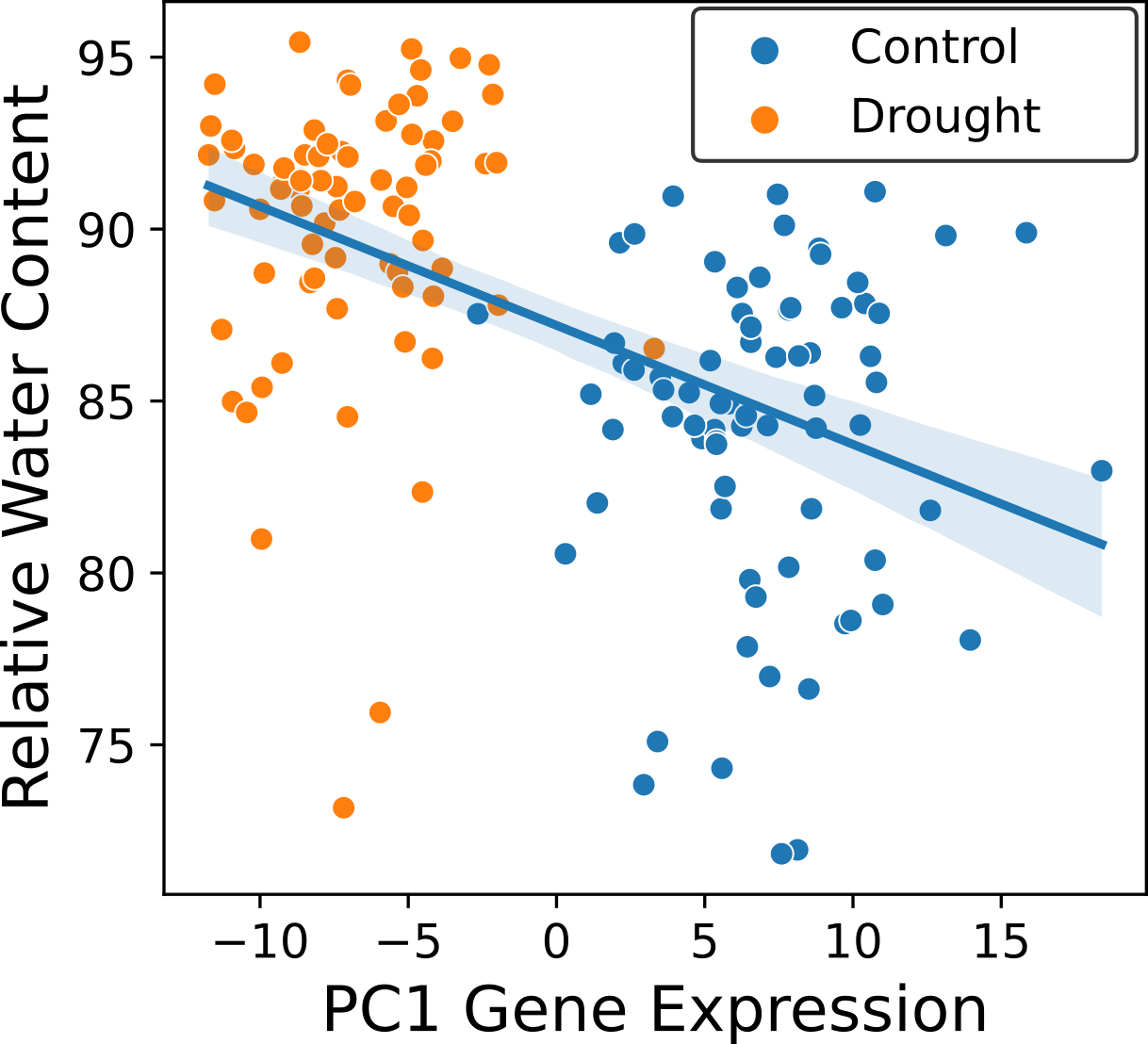

**Supplementary Figure 6.** First principle component of gene expression of top predictors in the sorghum trained model regressed against relative water content.

**Supplemental Tables**

**Supplemental Table 1. Summary of sorghum differential gene expression under drought in the field. (see external file)**

**Supplemental Table 2. Enriched Gene Ontology terms of upregulated genes under water deficit in sorghum. (see external file)**

**Supplemental Table 3. Enriched Gene Ontology terms of downregulated genes under water deficit in sorghum. (see external file)**

**Supplemental Table 4. Summary of sorghum data used in this study.**

| **Accession Number** | **Study Name** | **Citation** |
| --- | --- | --- |
| PRJNA527782 |  | (Varoquaux et al., 2019) |
| PRJNA556240 |  | (Azzouz-Olden et al., 2020) |
| PRJNA319738 |  | (Fracasso et al., 2016) |
| ERP010515 |  | (KATIYAR et al., 2015) |
| PRJNA585370 |  | (Abdel-Ghany et al., 2020) |
| PRJXXX | Sorghum Field | This study |
| PRJXXX | Sorghum Chamber | This study |
| PRJXXX | Maize NAM | This study |

S**upplemental Table 5. Model training scheme for random forest classification.**

| **Model Description** | **Training set** | **Testing set** |
| --- | --- | --- |
| Sorghum Field Dataset only | 75% of genotypes | 25% of genotypes |
| Cross sorghum dataset model | All but one sorghum dataset | Left out dataset |
| Sorghum trained cross species model | Each individual sorghum dataset as well as all sorghum datasets combined | Maize dataset |
| Maize trained cross species model | Maize dataset | Each individual sorghum dataset, as well as all sorghum datasets combined |
| Cross Developmental Stage model | All vegetative sorghum samples | All flowering / post flowering sorghum samples |

**Supplemental Table 6. Grid search parameters used to tune random forest models**

| **Parameter** | **Search Space** |
| --- | --- |
| n_estimators | 25 - 1000 in steps of 10 |
| max_features | [“auto”, “log2”, “sqrt”] |
| max_depth | 10-100 in steps of 11, None |
| min_samples_split | [2,5,10] |
| min_samples_leaf | [1,2,4] |
| bootstrap | [True,False] |
